# Effects of carbon-based additive and ventilation rate on nitrogen loss and microbial community during chicken manure composting

**DOI:** 10.1101/2020.02.19.956029

**Authors:** Ruixue Chang, Yanming Li, Qing Chen, Xiaoyan Gong, Zicheng Qi

## Abstract

Aerobic composting is a sustainable method for recycling of chicken manure, while its unsuitable porosity and carbon to nitrogen ratio limit the oxygen supply, which must result in high nitrogen loss because of anaerobic micro-zones in the materials. Treatments with five carbon-based additives and two ventilation rates (0.18 and 0.36 L·min^-1^·kg^-1^ DM) were set in chicken manure composting, to investigate their effects on biodegradation process, ammonia (NH_3_) emission, nitrogen loss, physiochemical properties and microbial community. The additives and ventilation rates influenced the CO_2_ production from the 2^nd^ week, meanwhile varied the physiochemical parameters all the process. No inhibitory effect on the maturity were observed in all treatments. With woody peat as additive, the NH_3_ emission amount and nitrogen loss rate were shown as 15.86 mg and 4.02 %, when compared with 31.08-80.13 mg and 24.26-34.24 % in other treatments. The high aeration rate increased the NH_3_ emission and nitrogen loss, which were varied with different additives. The T-RFLP results showed that the additives and the ventilation rates changed the microbial community, while the prominent microbial clones belonged to the class of *Bacilli* and *Clostridia* (in the phylum of Firmicutes), and *Alphaproteobacteria, Deltaproteobacteria* and *Gammaproteobacteria* (in the phylum of Proteobacteria). *Bacillus spp.* was observed to be the most dominant bacteria in all the composting stages and treatments. We concluded that woody peat could improve chicken manure composting more than other additives, especially on controlling nitrogen loss. 0.18 L·min^-1^·kg^-1^ DM was suitable for chicken manure composting with different additives.

## 1. Introduction

Due to the rapid development of chicken farms in China, the output of chicken manure has risen sharply in the past decades, which was nearly 102 million tons (dry weight) in 2016 (Jia et al., 2018). The over production and accumulation of untreated chicken manure has caused a series of environmental and social problems (Shi et al., 2018). Recycling the chicken manure to arable land as fertilizers has been recognized as a sustainable utilization method, for chicken manure has high concentration of macro and micro nutrients than other livestock manure. Aerobic composting could effectively convert the livestock manure into fertilizer or amendments used to improve soil fertility and promote plant growth (Hageman et al., 2018). While the low contents of lignocellulose and low carbon to nitrogen ratio of chicken manure would limit the oxygen consumption and organic matter degradation during chicken manure composting (Wang et al., 2015), which may contribute to more nitrogen loss and anaerobic microdomains. Different additives and suitable rate of forced ventilation used to improve the porosity and C/N ratio could make great significance to reduce the emission of GHGs and NH_3_, mitigate the mobility of heavy metals, and conserve other essential nutrients for chicken manure composting (Awasthi et al., 2017; Mao et al., 2018). Many researches have confirmed the effects of additives such as zeolite (Awasthi et al., 2016; Chan et al., 2016), bentonite (Wang et al., 2016), medical stone (Wang et al., 2017), woody peat and biochar (Zhang et al., 2014; Awasthi et al., 2017; Chang et al., 2019b), saw dust (Sharma et al., 2018), pine bark (Brito et al., 2015) and peanut hull (Erickson et al., 2014), for various organic waste composting to mitigate the emission of NH_3_ and GHGs and conserve the nutrients. However, the varied biodegradable organic matter content in these additives would significantly influence their improvement of composting process and the temperature (Chang et al., 2019a). Forced ventilation is used to supply adequate O_2_ during composting, which was found to result in lower GHG emissions when compared with physical turning and passive ventilation systems (Hao et al., 2001; Park et al., 2011). Most previous studies have reported that losses of NH_3_ increase and CH_4_ emissions decrease with increasing ventilation rate (Osada et al., 2000; Jiang et al., 2011; Shen et al., 2011). However, the values of N_2_O emissions in composting could not follow any trend when the increasing ventilation rate changes (Osada et al., 2000; Shen et al., 2011; Jiang et al., 2011). These inconsistent results may be caused by the different C and N dynamics and the O_2_ consumption in various raw materials because of their different physicochemical characteristics and biodegradable organic matter content. In the present study, considering the probable influence of carbon-based additives and ventilation rates on composting and the nutrient loss, we carried out 2 series of 30-d chicken manure composting in a lab-scale composting system, to explore the effects of carbon-based additives with different biodegradable organic matter, and the effects of high and low ventilation rates with the same additives on CO_2_ and NH_3_ emissions, composting process and the microbial community changes.

## 2 Material and methods

### 2.1 Set-up of experiment

Chicken manure, corn straw, saw dust, pine bark and peanut hull were collected from local greenhouse and farmland in Beijing, China. The additives (corn straw, saw dust, pine bark, peanut hull) were air dried and cut into 2-3 cm pieces to obtain a uniform particle size that enabled good mixing. Woody peat was supplied by View Sino international Ltd., which was used in powder form. Main characteristics of the raw materials were shown in Table 1.

**Table 1.** Physical and chemical properties of the materials

| Materials | Total carbon content / % | Total nitrogen content / % | C/N | Moisture / % | Cellulose / % | Lignin / % |
| --- | --- | --- | --- | --- | --- | --- |
| Chicken manure | 37.63 | 4.33 | 8.69 | 81.28 | 13.06 | 9.06 |
| Wheat straw | 62.35 | 0.73 | 85.41 | 7.14 | 42.03 | 8.86 |
| Woody peat | 62.84 | 0.59 | 196.51 | 14.04 | 2.92 | 28.91 |
| Saw dust | 67.70 | 0.41 | 165.12 | 8.33 | 50.85 | 17.39 |
| Pine bark | 73.52 | 0.44 | 167.09 | 8.79 | 61.62 | 31.22 |
| Peanut hull | 52.77 | 1.06 | 49.78 | 7.85 | 64.10 | 25.65 |

The experiments were conducted in a bench-scale composting system (Fig. 1) in the lab of China Agricultural University, designed to simulate the temperature changing without external effects (e.g., heat loss) (Michel and Reddy, 1998; Meng et al., 2016). The system details were described in our previous study (Chang et al., 2019a). There were two series of treatments in our experiments, whose materials ratios were shown in Table 2.

**Table 2.**
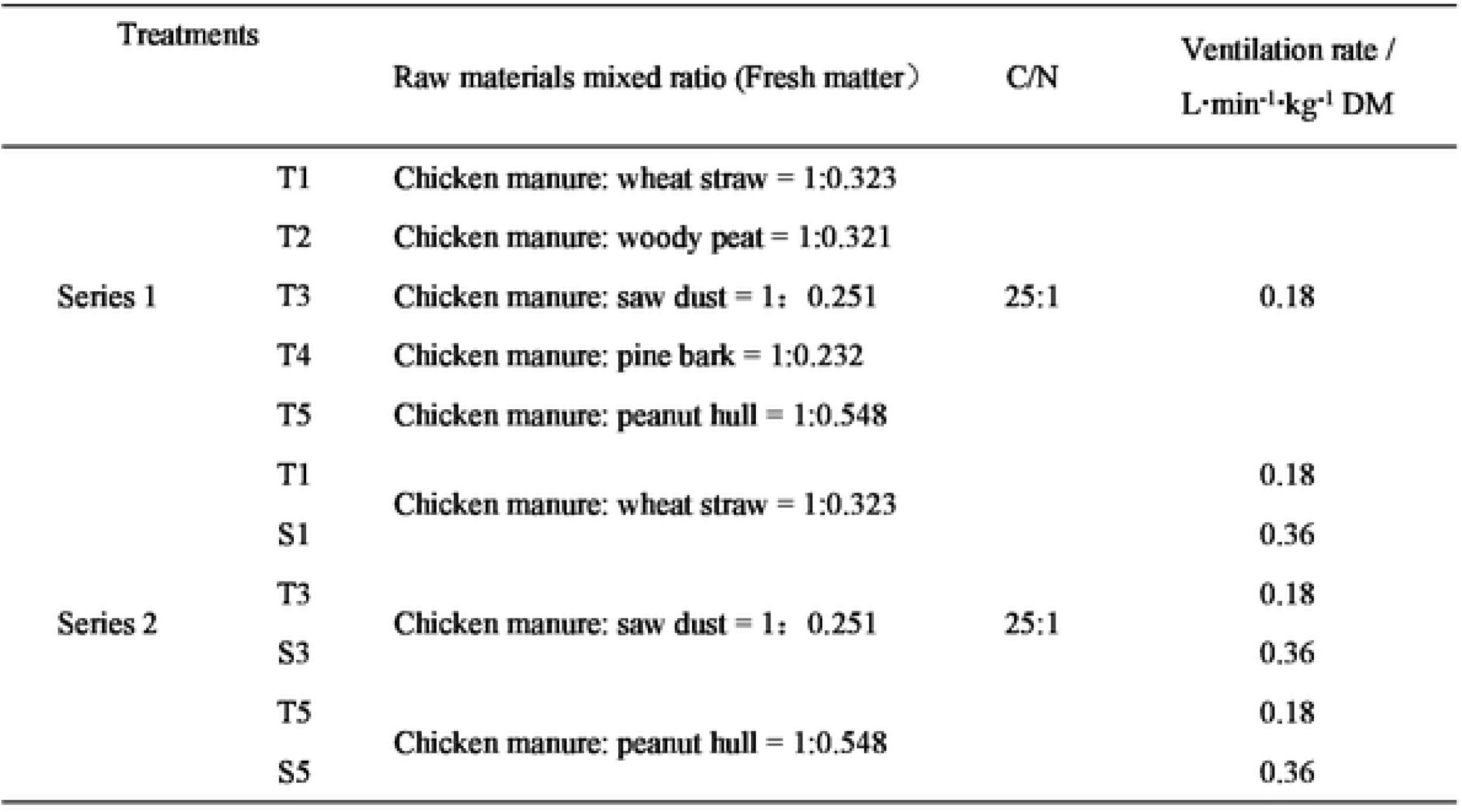
Experiment design of different resource carbon

**Fig 1.**
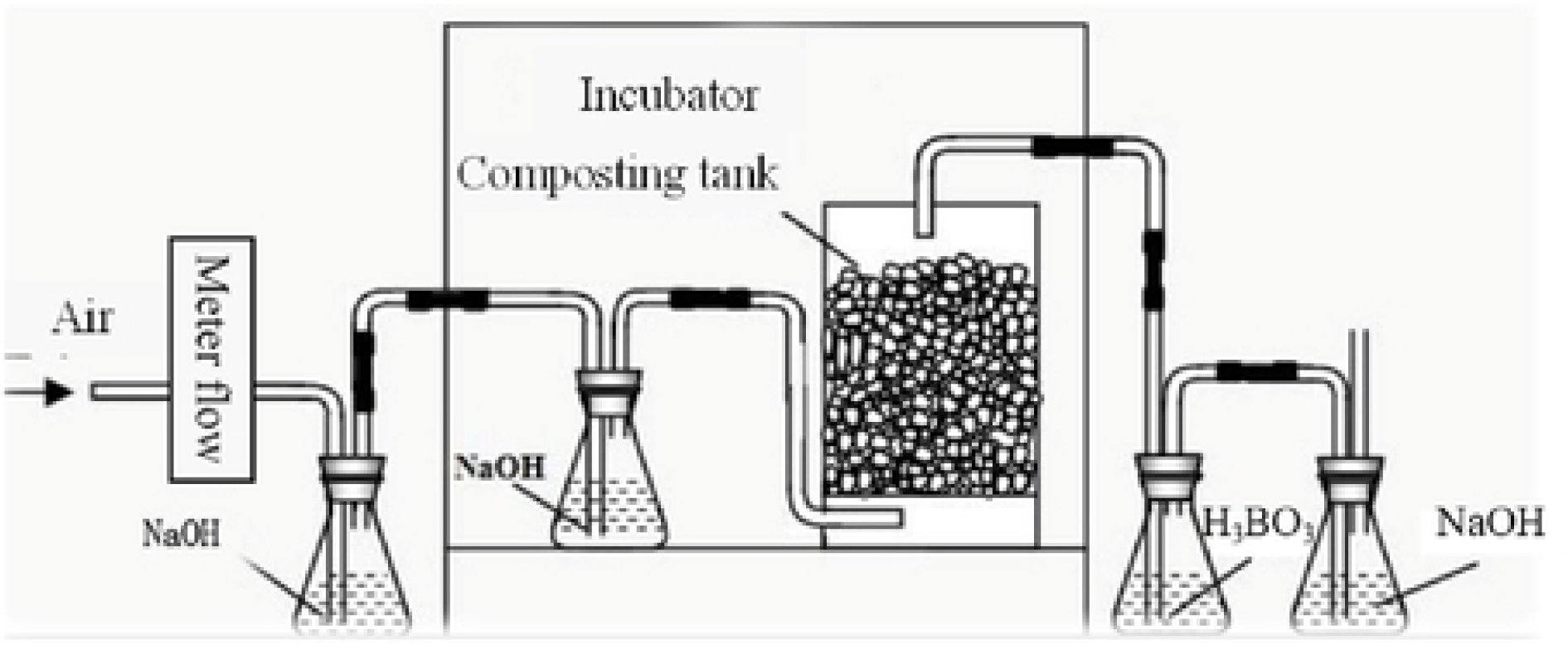
Diagram of composting system.

### 2.2 Samples collection and analysis

During the process, solid samples were collected on the days of 0, 3, 7, 14, 21, 28 and 35 after mixing well. Each sample was thoroughly mixed and then divided into two parts: one part was air-dried to analyze physicochemical characteristics, like total nitrogen (TN) and ash content; the other part was stored in the freezer at −20 °C for determination of other parameters, like pH value, Electric Conductivity (EC), Germination Index (GI), extractable ammonium and microbial community. A 1:5 aqueous extract (w/v) of the fresh composts with 2N KCl solution was used for the analysis of extractable ammonium (NH_4_ ^+^-N), and NH_4_ ^+^-N was analyzed by SEAL Analytical (BL-TECH). Measurement of other parameters and calculation methods were followed the methods shown in Chang et al. (2019a and 2019b).

### 2.3 Analysis of microbial community

Total community DNA was extracted from 0.5 g compost samples using the FastPrep DNA kit (MP Biomedicals, Santa Ana, CA) according to the manufacturer’s protocol (Feng et al, 2012). The extracted DNA solutions were diluted in suitable times. The 16S rRNA genes were amplified using universal bacterial primers: 27f forward (5’-AGAGTTTGATCCTGGCTCAG-3’) and 907r (5’-CCGTCAATTCMTTTGAGTT −3’) reverse. The 27f forward primer was labeled with 6-carboxyfluorescein (FAM). Each PCR reaction mixture contained 50 µl liquid: 37.5 µl dd H_2_O, 10* PCR reaction buffer 5µl (Tiangen Biotech, Beijing), dTNPs 4µl, 27f-FAM 0.75µl, 907r 1.5 µl, BSA 0.5µl, rTaq DNA polymerase0.5 µl (TakaRa), DNA template 2µl. The reaction mixture was incubated at 94°C for 4 min, and then cycled 30 times through three steps: denaturing (94 °C; 45 s), annealing (52°C; 45 s), and primer extension (72°C; 60 s) in a PTC-100 thermal cycler. Then the last step is 10 mins’ primer extension. Amplification product sizes were verified by electrophoresis in 2.0% agarose and ethidium bromide staining. To obtain sufficient DNA for T-RFLP analysis and to minimize PCR bias, amplicons from three PCR runs for each root sample were combined (Clement et al., 1998) and then purified using a PCR purification kit (PCR Clean-up Kit; PROMEGA Inc., Wisconsin, USA).

To construct bacterial 16S rRNA gene-based clone libraries, we prepared DNA samples extracted from five compost samples with the richest bacteria diversity from different additive treatments, respectively. The PCR amplification used the same primers as those indicated above. PCR products were purified and ligated into the pMD19-T Vector (TakaRa) according to the manufacturer’s instructions. 1 ml suction head was used to blow and absorb the bacteria at the bottom of the centrifuge, and then 20-40 µl of the cells were coated on LB AGAR plate medium containing X-Gal, IPTG and Amp for overnight culture at 37°C, to form a single colony. White clones were selected and underlined on LB-Amp plates, and cultured overnight at 37°C. The screened positive clones were sent to the sequencing company for sequencing, which were screened with the primers M13-47 (5’-CAGCAC TGA CCC TTT TGG GAC CGC-3’) and RV-M (5’GAG CGG ATA ACA ATT TCA CAC AGG-3’). Enter the results into NCBI GeneBank database and perform Blast search to obtain similar gene sequences. Phylogenetic tree was constructed with MEGA software and NJ method (neighbor-joining).

### 2.4 Statistical analysis

All the results were summarized and figured in Excel. Statistical comparisons were performed using SPSS v.18.0 software with the two-way ANOVA analysis of variance test. A probability was defined with a least significant difference at two sides of P < 0.05.

## 3. Results and discussion

### 3.1 Biodegradation estimated by accumulative amount of CO_2_

Rapid decrease of total organic carbon and increase of cumulative CO_2_ amount coincided with the biodegradation of organic matter and the rise of temperature during composting (Wong and Fang, 2000). As shown in Fig. 2A, similar CO_2_ emission amount were observed in the first 7 days when the ventilation rate was 0.18 L·min^-1^·kg^-1^ DM, suggested the easily-degraded organic matter was degraded and transferred to CO_2_. Then the decreased rates in T2-T5 indicated that the easily-degraded organic matter were less in these treatments than T1. For the concentrations of cellulose and lignin were higher in saw dust, pine bark and peanut hull, when compared with wheat straw, while cellulose and lignin were hard to be biodegraded directly (Chang et al., 2019b). Woody peat is rich in carbon and humus but unavailable for microbes, so that the CO_2_ amount was lower than that in T1. Similar result was observed in Chang et al. (2019b), in which woody peat and corn stalk were used in vegetable wastes or sewage sludge composting. When the ventilation rate was increased to 0.36 L·min^-1^·kg^-1^ DM, less CO_2_ emission amount were observed in S1, S3 and S5(Fig. 2B), suggested that the ventilation rate of 0.18 L·min^-1^·kg^-1^ corresponded to a higher biodegradation than 0.36 L·min^-1^·kg^-1^ DM in the current study. Significantly differences were observed after the first 7 days. As shown in Qasim et al. (2019), in the ventilation range of 0.3-0.9 L·min^-1^·kg^-1^ DM when composting with poultry manure and sawdust, the low aeration rate (0.3 L·min^-1^·kg^-1^ DM) corresponded to a higher and longer thermophilic phase than did the high aeration rate (0.9 L·min^-1^·kg^-1^ DM). The CO_2_ volatilization was directly related to the temperature profile of the substrate, shown as significant differences in the 2^nd^ and 3^rd^ weeks of composting but none in 1^st^ week. What’s more, several previous studies have recommended the aeration methods and rates as, 0.44 L·min^-1^·kg^-1^ DM in the composting of maize stalks and cow feces (Nada, 2015), 0.62 L·min^-1^·kg^-1^ volatile solids (VS) in the composting of vegetable and fruit wastes (Arslan et al., 2011), 0.5 L·min^-1^·kg^-1^ DM in the composting of chicken manure and sawdust (Gao et al., 2010), 0.25 L·min^-1^·kg^-1^ DM in the composting of dairy manure with rice straw (Li et al., 2008), 0.43-0.86 L·min^-1^·kg^-1^ DM in the composting of food waste (Lu et al., 2001), etc. All of these suggested that the aeration rate should be set according to the compost material and composting process, based on the oxygen needed and supplied during the process.

**Fig. 2.**
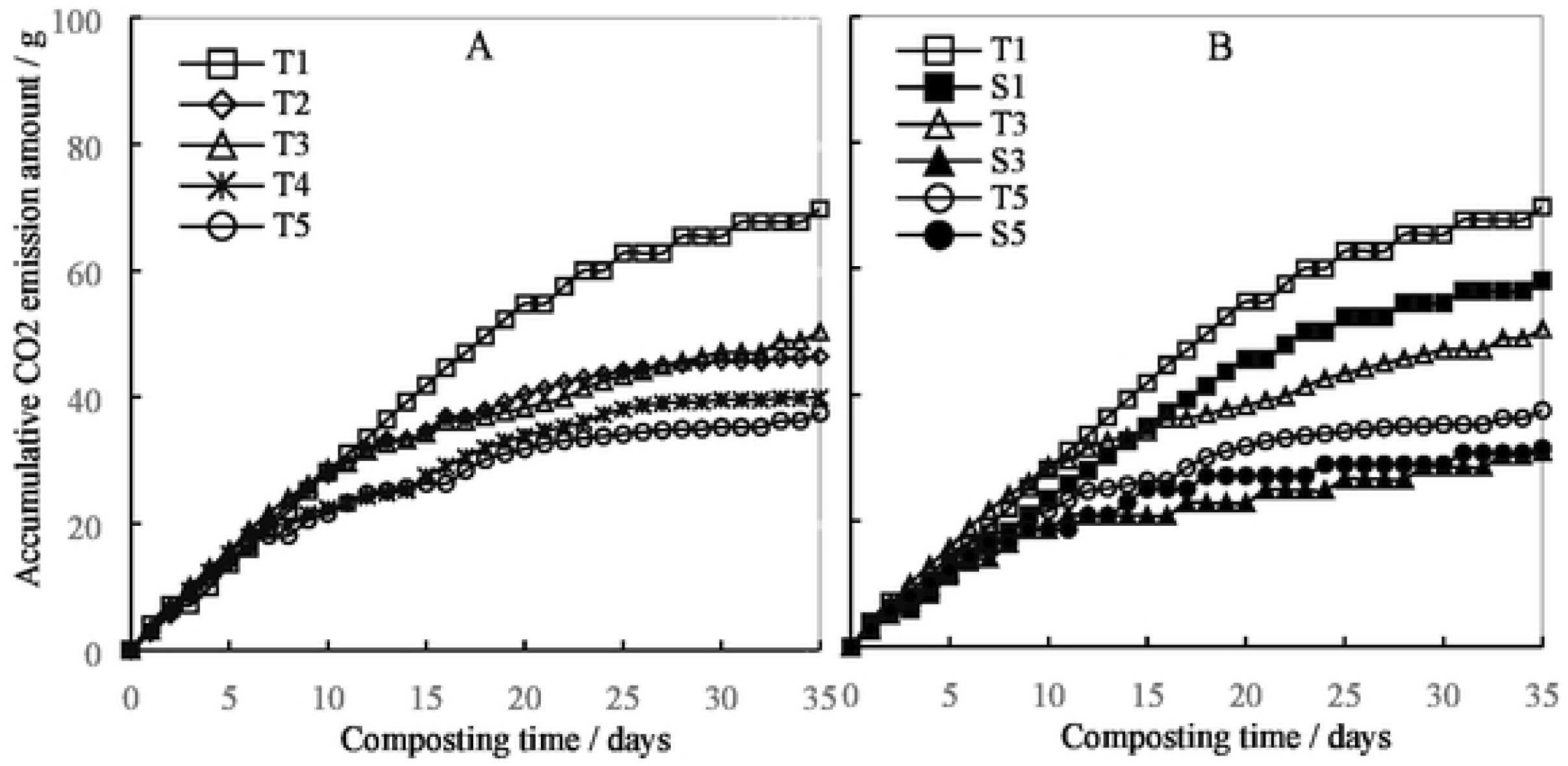
Effects of carbon-based additives and ventilation rate on CO_2_ emissions during chicken manure composting.

### 3.2 Physiochemical characteristics

The appropriate pH range for maintaining high microbial activity during composting is 7-8, which would be changed along with the biodegradation of organic matter. For the complex components were degraded to organic acids and then to CO_2_, meanwhile CO_2_, NH_3_, other gases and volatile organic acids were emitted from the composting system (Eklind and Kirchmann, 2000). As shown in Fig.3A, the pH values in all the treatments were in the range of 6.8∼8.4, suggested the carbon additives made no difference on the biodegradation process. Even the additives used in current experiment changed pH value of the products, the final value were all in the range of 7.0∼8.2, which is good for agricultural utilization (Maso and Blasi., 2008). The pH value in T2 was lower than others, indicated the potential advantage of woody peat, to reduce the NH_3_ emission by decreasing the material pH value. Increase of the aeration rate quickly increased the pH values (Fig. 3B), for the gases and volatile organic acids were forced to emitted more frequently than in the low aeration rate treatments. Then the reduction of biodegradation resulted in stable change of pH value, similar as those shown in the low aeration rate treatments. The final pH values of products in S1-S3 were higher than those in T1-T3.

**Fig. 3.**
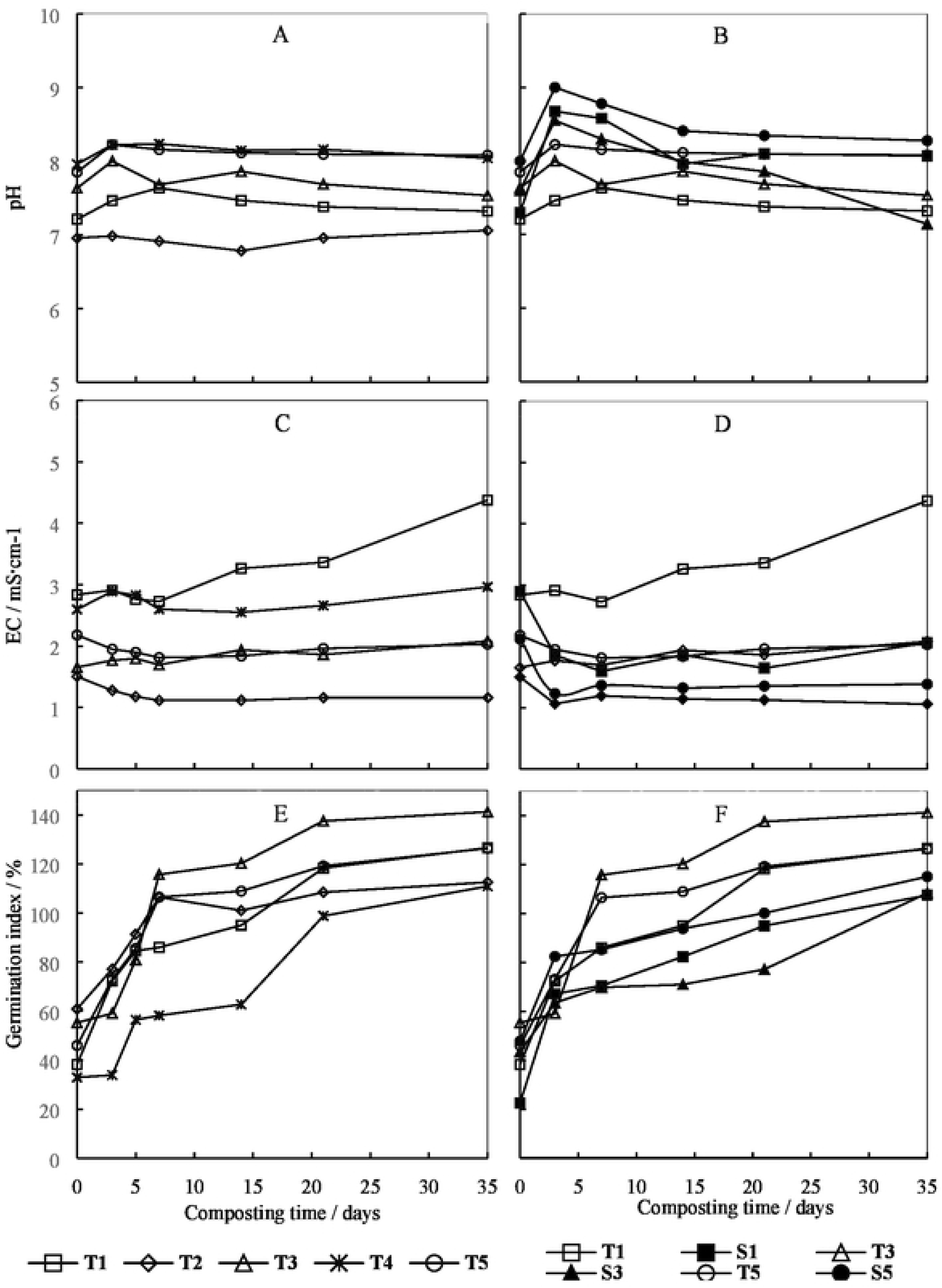
Effects of carbon-based additives and ventilation rate on physiochemical characteristics during chicken manure composting

For the compost products are always used as organic amendments or organic fertilizer in soil (Liu et al., 2011), EC should be under 4 mS·cm^-1^, which reflects no inhibitory effects on plant growth from the compost products (Li et al., 2007). A slightly decrease was shown in the first several days for almost every treatment, followed with a stable value till the end (Fig. 3C and 3D). The carbon additives influenced the EC variation, while they were always in the safe range, except 4.37 mS·cm^-1^ in T1. For more biodegradation happened in T1, which was indicated by the CO_2_ production and temperature. Woody peat used in T2 reduced the EC in the whole process, because of the absorption caused by its rich humic acid. The rapid emission of gases and volatile organic acids in treatments with high aeration rate also reduced the EC value caused.

To avoid the toxic effects on plant growth resulted from the toxic substances, such as short-chain fatty acids, GI is always used as an important index to evaluate whether compost is mature enough. A minimum value of 80% is considered to indicate the compost mature at an extraction ratio of 1:5 (compost: water wet w/v). As shown in Fig. 3E and Fig. 3F, the GIs increased with the decomposition of toxic materials, especially in the 1^st^ week. Nearly all the GIs were higher than 80% in the treatments, except T4 in series I, and S1 and S3 in series II. Then the GIs keep slightly increasing till the end of the process, with the GIs over 100%. The results indicated that the five organic wastes chose in the current study all could used to composted with chicken manure, by adjusting the free air space and C/N. While the higher aeration rate (0.36 L·min^-1^·kg^-1^ DM) decreased the maturity process. However, it was opposite in a pig manure composting from Guo et al. (2012), the suitable aeration was 0.48 L·min^-1^·kg^-1^ DM when considering the maturity (Guo et al., 2012), even the lower aeration rate (0.24 L·min^-1^·kg^-1^ DM) had better performance on biodegradation. The reasons should be complex, one is the aeration was intermittent in Guo et al., (2012) so that higher aeration rate could supply enough O_2_ than lower one. The other is that lower C/N of the mixed materials (< 20) make high concentration of TAN (total ammonium nitrogen) in the materials, which would inhibit the seedling.

### 3.3 NH_3_ emission and nitrogen concentrations

The changes of accumulative NH_3_ emission amount during the composting of chicken manure with different additives and different ventilation rates were shown in Fig. 4A and Fig. 4B. Generally, the accumulative amounts rapidly increased from the beginning to the 10^th^ day in T1-T3, while to the 20^th^ day in T4 and T5. Then the amount increased slowly till the end of the process. The significantly lower cumulative NH_3_ emission in T2 (15.86 mg) was related to the characteristics of woody peat, than those with straw, which were widely used as additives during manure composting. Low pH and rich of humic acid contributed to the absorption of NH_4_^+^, which was proved by previous studies (Chang et al., 2019b). More cumulative NH_3_ amounts were observed in T3-T5 than that in T1, which may be related with the lower biodegradable organic carbon in the mixed materials. For there are high concentrations of lignocellulose in saw dust, pine bark and peanut hull, which may decrease the biodegradable C/N and increase the NH_3_ emission (Chang et al., 2019b). Comparing treatments with the same compost materials, S1 had significantly higher cumulative NH_3_ losses (by 117.70%) compared with T1, S3 (by 25.55%) with T3, and S5 (by 38.14%) with T5. Which suggested the increased aeration rate led to higher cumulative NH_3_ losses.

**Fig. 4.**
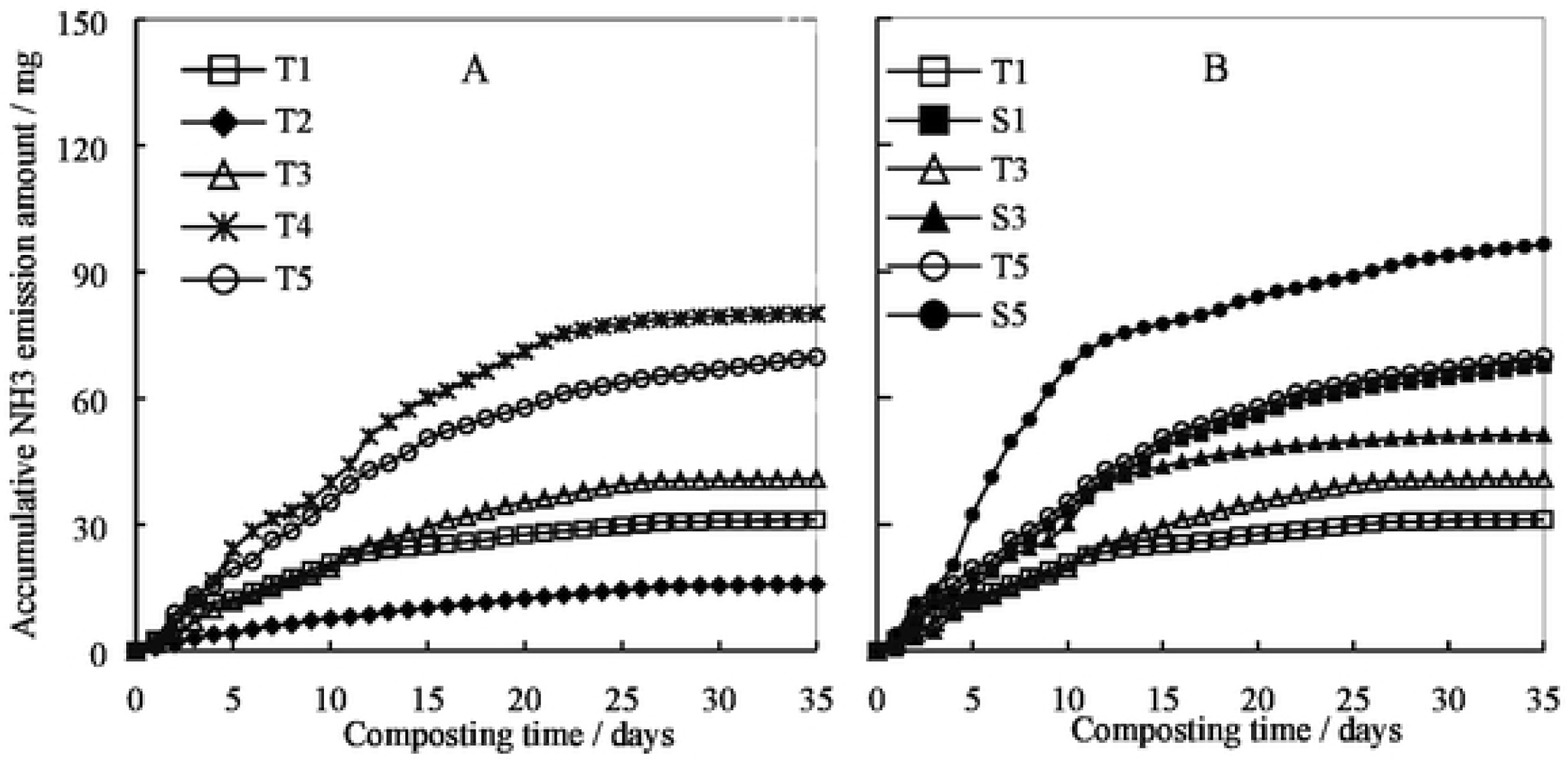
Effects of carbon-based additives and ventilation rate on NH_3_ emissions during chicken manure composting.

During the composting, the TN always increase from the initial to the end, because of the concentration effect caused by the significant organic decomposition (Chan et al., 2016). In current study, most of the treatments were consistent with this theory, except T5 and S5, in which the peanut hull was used as carbon additives (Table 3). For their NH_3_ emission were really high when compared with other treatments, shown as 69.79 mg and 96.41 mg (Table 3). What’s more, less matter loss resulted from less organic decomposition decreased the concentration effect, for a high ratio of peanut hull was mixed with the chicken manure, which contained high concentrations of cellulose and lignin. There are several forms of nitrogen in the compost, among which organic nitrogen (Nor) and nitrate nitrogen (NO_3_^-^-N) are normally stable in the materials and called stable nitrogen (Chang et al., 2019b), while NH_4_ ^+^-N may transformed to NO_3_^-^-N or emitted as NH_3_, when the temperature was decreased or during the utilization of the compost in arable land. High NH_4_^+^-N concentration in the materials always contributes to high NH_3_ emission rate, which was consistent in our study except T2. The different results indicated the important function of woody peat to absorb and fix the NH_4_^+^-N, because of its porous structure and rich of humus acid (Chang et al., 2019b). Similar results could be observed when biochar was used during composting (Qiu et al., 2019). More NH_4_^+^-N in treatments T3 and T5 than S3 and S5 (shown in Table 3), suggested high aeration rate helped to transfer NH_4_^+^-N into NH_3_ ventilation, so that more NH_3_ ventilation were observed in Fig.5 and Table 2. Meanwhile the total nitrogen loss rates were higher in treatments with high aeration rate resulted from high NH_3_ ventilation.

**Fig. 5.**
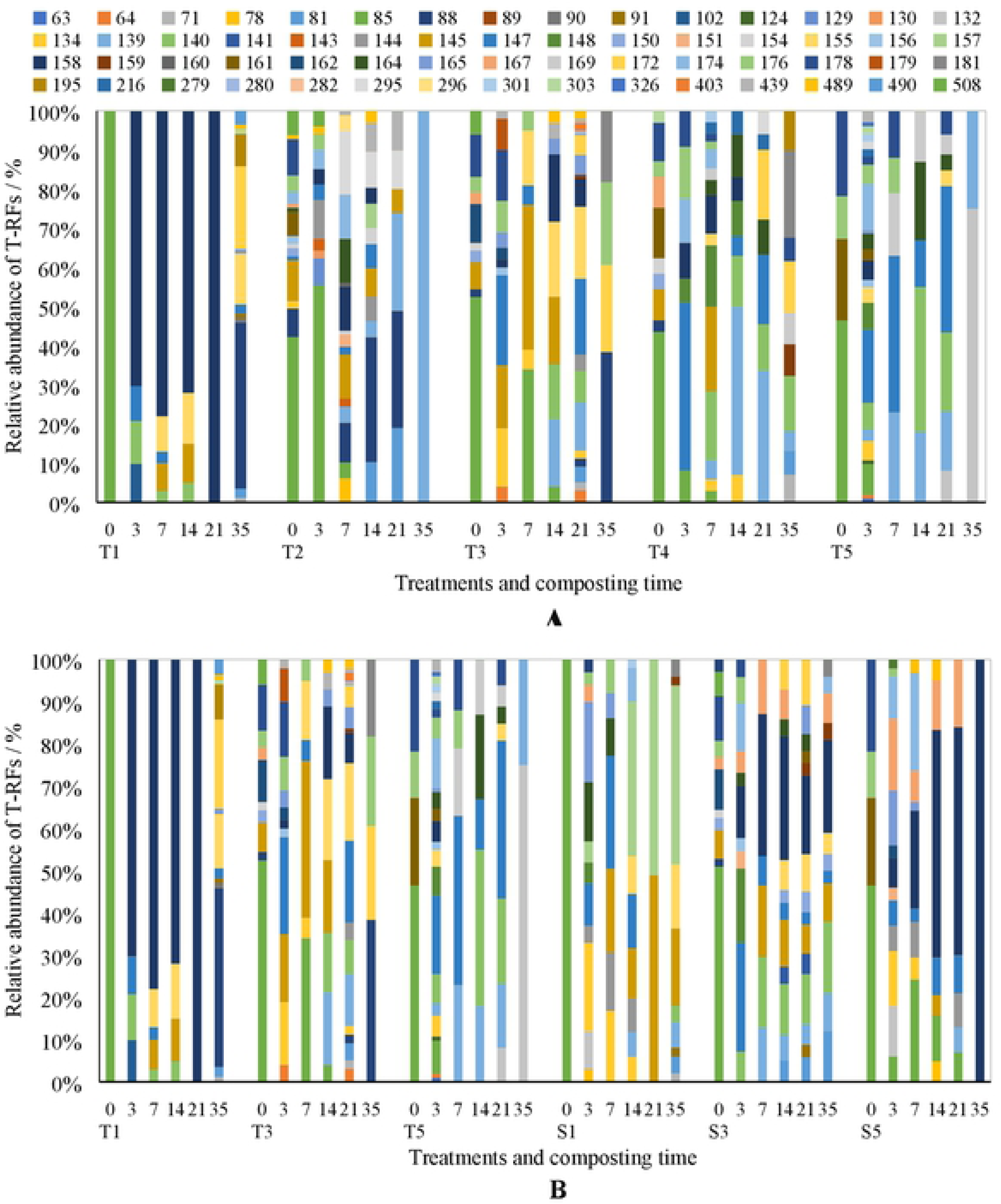
Effects of carbon-based additives (A) and ventilation rate (B) on relative abundance of T-RFs during chicken manure composting.

**Table 3.** Nitrogen transformation and nitrogen loss in all treatments

|  | Treatments | TN / g·kg<sup>-1</sup> |  | Ammonium nitrogen / mg·kg<sup>-1</sup> |  | Total NH<sub>3</sub> emission amount / mg | TN loss / % |
| --- | --- | --- | --- | --- | --- | --- | --- |
|  |  | Initial | End | Initial | End |  |  |
| Series 1 | T1 | 21.55 | 23.52 | 759.07 | 105.40 | 31.08 | 24.26 |
|  | T2 | 19.03 | 21.60 | 1310.33 | 840.73 | 15.86 | 4.02 |
|  | T3 | 23.89 | 26.44 | 963.93 | 212.07 | 40.79 | 28.45 |
|  | T4 | 19.85 | 20.65 | 1632.33 | 817.20 | 80.13 | 34.24 |
|  | T5 | 20.88 | 16.68 | 1236.73 | 877.80 | 69.79 | 32.74 |
| Series 2 | T1 | 21.55 | 23.52 | 759.07 | 105.40 | 31.08 | 24.26 |
|  | S1 | 21.55 | 22.19 | 759.07 | 138.80 | 67.66 | 35.75 |
|  | T3 | 23.89 | 26.44 | 963.93 | 212.07 | 40.79 | 28.45 |
|  | S3 | 23.68 | 25.06 | 963.93 | 154.80 | 51.21 | 29.51 |
|  | T5 | 20.88 | 16.68 | 1236.73 | 877.80 | 69.79 | 32.74 |
|  | S5 | 20.88 | 16.73 | 1236.73 | 494.30 | 96.41 | 36.24 |

### 3.4 Structure of microbial community

Compared with the normal additive corn straw. woody peat, saw dust and pine bark increased the biodiversity at the beginning while peanut hull increased it at the 3^rd^ day (Fig. 5(A)). At the beginning, the most abundant T-RF was 85bp, with the ratio of 100%, 56%, 53%, 44% and 47%, in T1-T5 respectively. The most abundant T-RFs were shown in the 3^rd^ day of T2 and T4, while 7^th^ day of T3 and T5. After then, the numbers decreased significantly. In all the 60 T-RFs shown in the figures, 139bp, 145bp, 147bp, 174bp, 176bp and 178bp were shown high relative abundance in two or more treatments in different stages, indicated these T-RFs should be related with the biodegradation of organic matter. While 158bp in T1, 295bp in T2, 155bp in T3, 140bp in T4 and 132bp in T5 were shown high relative abundance in only this treatment, which should be a signal that this T-RF should be specific for this additive only. The results suggested the additives in the current study influenced the microbial community and population, which would result in different biodegradation process. The increase of the ventilation rate contributed to the higher biodiversity in all the three treatments (Fig. 6(B)), even it occurred in the first 7 days in S1 and S5 while in later two weeks in S3.

**Fig. 6.**
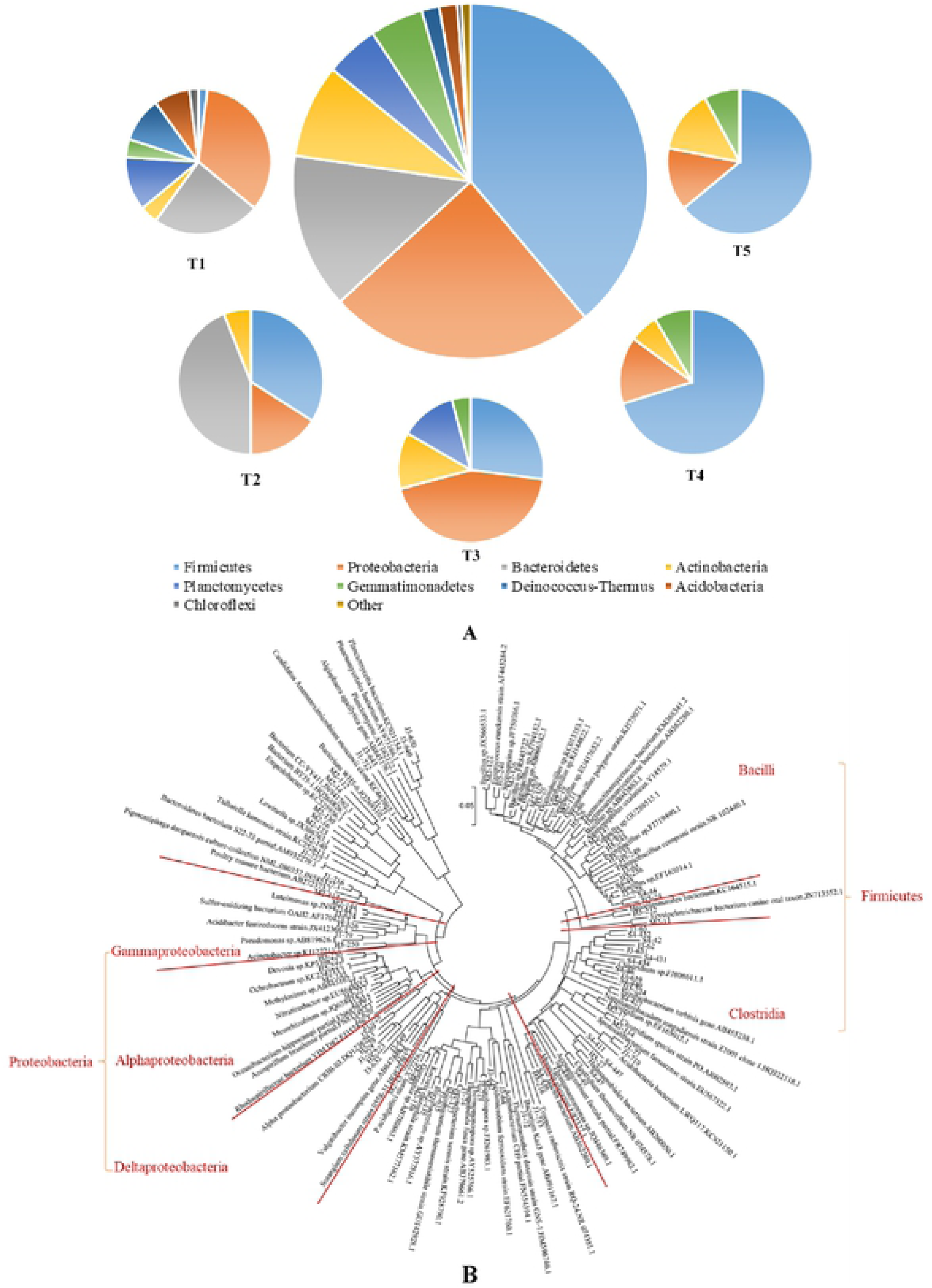
Composition of 247 sequences in 5 compost samples based on the bacteria clone library analysis (A) and their phylogenetic analysis (shown as OUT and restriction enzyme cutting site) (B)

The species distribution of bacteria in composting system can be better understood by constructing bacterial clone library. In this experiment, five of the compost samples in different treatments were selected to construct the clone library. Library analysis showed that the 247 sequences belonged to 9 different phyla, among which Firmicutes, Proteobacteria, Bacteroides and Actinomycetes account for 85.83% in the total sequences (Fig. 6(A)). Nearly 70 of the 247 sequences were Bacillus spp., which belongs to the phylum of Firmicutes. Their character of high-temperature resistance made it important on organic matter biodegradation during composting (Koyama et al., 2018). The result of the phylogenetic analysis affiliated with uncultured groups using the neighbor-joining method were shown in Fig. 6(B). Prominent clones belonged to an uncultured group in the phylum Firmicutes (class of *Bacilli* and *Clostridia*) and Proteobacteria (class of *Alphaproteobacteria, Deltaproteobacteria* and *Gammaproteobacteria*). Carbon additives used in different treatments influenced the clones, which were consistent in both Fig. 6(A) and Fig. 6(B). As Qiu et al. (2019) indicated, the kind of manure and biochar addition would both influence the bacteria community, especially when the materials were composted different duration. While it is similar that Firmicutes (class of *Bacilli* and *Clostridia*) and Proteobacteria (class of *Alphaproteobacteria, Deltaproteobacteria and Gammaproteobacteria*) were the main categories in the later time of composting. High proportions of Actinobacteria in the samples of T1 and T3 suggested that the addition of corn straw or saw dust may facilitate the growth of Actinobacteria and accelerate the degradation of lignocelluloses during the maturity stage, similar results were observed and proved in Qiu et al. (2019).

## 4. Conclusion

The biodegradation process and the pH, EC and GI values were influenced by the five carbon-based additives used in our experiment, while no inhibitory on composting maturity were observed. The aeration rate of 0.18 L·min^-1^·kg^-1^ DM was more suitable than 0.36 L·min^-1^·kg^-1^ DM for chicken manure composting. Woody peat had shown better effect on reducing NH_3_ emission and nitrogen loss, while more NH_3_ emission and nitrogen loss were observed when the aeration rate was higher. The prominent clones of the compost samples belonged to the phylum Firmicutes (class of *Bacilli* and *Clostridia*) and Proteobacteria (class of *Alphaproteobacteria, Deltaproteobacteria* and *Gammaproteobacteria*). Carbon-based additives and ventilation rates set in our experiment had made influences on the microbial community, while Bacillus spp. was always the most important one. Therefore, woody peat could be used as carbon-based additives instead of corn straw, and the suitable ventilation rates could reduce the NH_3_ emission and nitrogen loss. Additives and ventilation rates would not influence the prominent bacteria that used to promote the composting process.

## Acknowledgement

Financial supported was provided by National Key R&D Program of China (2018YFC1901000), China Agriculture Research System (CAR-23-B16) and Shandong Academy of Agricultural Sciences “Science and Technology Innovation Project” (CXGC2018D06)

